# SpatialPPI: three-dimensional space protein-protein interaction prediction with AlphaFold Multimer

**DOI:** 10.1101/2023.12.14.571766

**Authors:** Wenxing Hu, Masahito Ohue

## Abstract

The rapid advancement of protein sequencing technology has resulted in a gap between proteins with identified sequences and those with mapped structures. Although sequence-based predictions offer insights, they can be incomplete due to the absence of structural details. On the other hand, structure-based methods face challenges with newly sequenced proteins. AlphaFold emerges as a potential solution, especially in predicting protein-protein interactions. This research delves into using deep neural networks, specifically analyzing protein complex structures as predicted by AlphaFold Multimer. By transforming atomic coordinates and utilizing sophisticated image processing techniques, the study extracts vital 3D structural details from protein complexes. Recognizing the significance of evaluating residue distances in protein interactions, the study leverages image recognition approaches, notably integrating DenseNet and ResNet within 3D convolutional networks for protein 3D structure analysis. When benchmarked against leading protein-protein interaction prediction methods like SpeedPPI, D-script, DeepTrio, and PEPPI, our proposed method, named SpatialPPI, exhibits notable efficacy, emphasizing the promising role of 3D spatial processing in advancing the realm of structural biology.

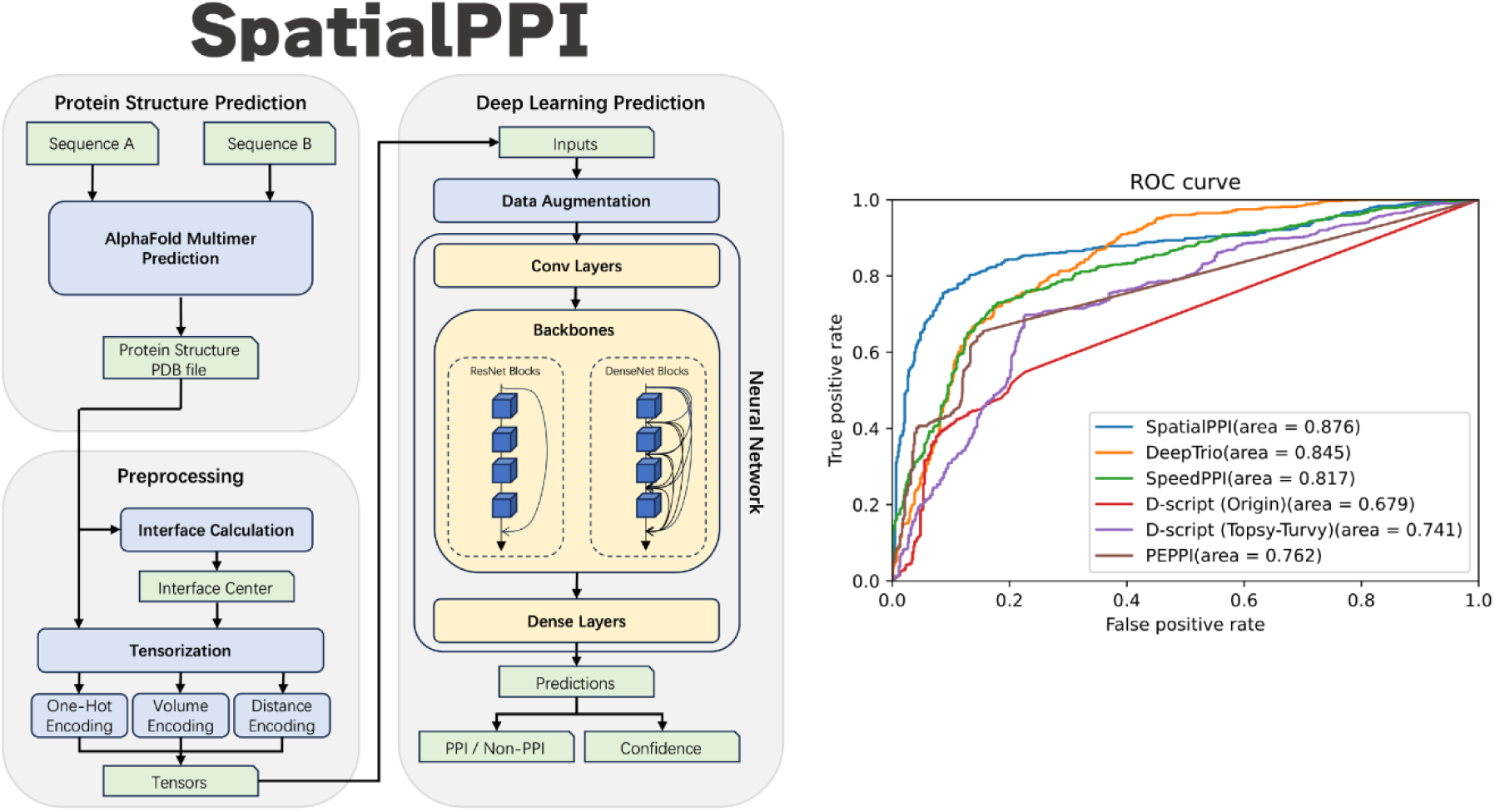

## 1. Introduction

Proteins are important building blocks of living organisms and carry out various biological functions, including catalyzing chemical reactions, transporting molecules, and serving as structural components of cells and tissues. The prediction of protein-protein interactions (PPIs) is a challenging task because proteins interact with each other in a highly complex and dynamic manner. Accurate prediction of PPIs can provide insights into the molecular mechanisms underlying biological processes and can assist in the development of new therapeutic interventions for diseases. Currently, the dominant approaches to predict PPIs include sequence-based prediction [1–4], which uses the amino acid chains of two proteins for the input, and structure-based predictions [5–6], which uses experimentally determined three-dimensional structures. Furthermore, the PPI predictions also can be categorized into similarity methods based on homologous interactions [7–8], and machine learning based methods [9–12].

The exponential growth rate of known protein sequences, promoted by the rapid development of automated sequencing technologies, has led to a significant disparity in the pace of growth between protein sequence determination and protein structure experimental determination. This trend has been widely noticed [13], and is primarily due to the inherent complexity and time-consuming nature of experimental methods used to determine protein structures. On the other hand, determining the sequence of a protein is relatively straightforward and can be achieved using automated sequencing technologies. There was an exponential sequence-structure gap found between the number of known structures in Protein Data Bank (PDB) [14] and the number of known sequences in UniProt [15]. However, predicting PPIs based on three-dimensional structure is advantageous over sequence-based prediction because PPIs are frequently mediated by specific structural features, such as complementary binding surfaces and specific hydrogen bonds, which can be more accurately modeled using 3D structure information than sequence information alone [16]. In contrast, sequence-based predictions of PPIs rely on identifying conserved motifs and domains that are involved in interactions, which can be at a disadvantage due to the diversity of potential binding partners and the limited information provided by sequence alone [17]. Furthermore, 3D structure-based predictions of PPIs offer a more comprehensive view of the interactions, including the types of interactions, such as hydrophobic interactions and electrostatic interactions, and the specific residues involved [18]. This detailed information can provide insights into the mechanisms underlying PPIs, which can inform drug discovery efforts and the design of novel therapeutics. Overall, the use of 3D structure information for PPI prediction holds great promise for advancing our understanding of complex biological processes and developing new treatments for diseases. However, due to the complexity of obtaining protein structure, the prediction method based on protein structure will face the problems of the insufficient amount of data and difficulty in predicting new proteins due to the period of the experiments.

The AlphaFold Multimer [19] could be a potential solution to the deficiency of structure-based PPI prediction methods. The AlphaFold Multimer is a computational tool that utilizes deep learning algorithms to predict the 3D structures of protein complexes, which offers a novel approach to investigating PPIs. Due to the complex nature of PPIs, predicting their structures and functions remains a challenge in the field of structural biology. However, the ability of AlphaFold Multimer to generate accurate and reliable predictions of protein complex structures could significantly enhance the understanding of the underlying mechanisms of these interactions. With the increased accuracy of structural predictions, insight could be gained into the binding sites and key residues involved in PPIs, thereby improving the ability to develop deep learning methods that predict these interactions.

Recent studies have introduced various novel methods to predict PPIs. One such approach is the FoldDock algorithm [20], which utilizes multiple sequence alignments (MSAs) generated by AlphaFold to pDockQ scores. The pDockQ score of FoldDock is calculated by a sigmoidal fit from the average pLDDT score [21] generated by AlphaFold. By doing so, the FoldDock algorithm can predict the potential for the two proteins to interact with each other and has achieved state-of-the-art accuracy. Another approach is PEPPI [22], which combines homologous search and a multi-layer perceptron classifier [23] using Gaussian kernel density estimation. Furthermore, Jones *et al*. [24] have proposed an approach that utilizes a neural network for transforming three-dimensional protein-ligand complexes into a 3D data structure by rescaling atomic coordinates at a 1 Å resolution to forecast protein-ligand binding affinity, which discussed a method for inputting protein structures to convolutional neural networks. In addition to these methods, three-dimensional convolutional neural networks (3D-CNNs) have found extensive use in computer vision domains such as medical diagnosis for 3D image segmentation [25–26] and recognizing gestures and activities in videos [27–28]. Together, these approaches provide a promising path for understanding complex interactions between proteins and improving the ability to predict and model their behavior.

In light of this, we propose SpatialPPI, which uses deep neural networks to analyze the protein complexes predicted by AlphaFold Multimer to predict PPIs. The spatial models of protein complexes are determined by transforming atomic coordinates by calculating atomic distributions. This approach employs advanced image processing strategies to extract crucial 3D structural information from the protein complexes. DenseNet[29] and ResNet[30] are network architectures that are fruitful in image recognition, and the applications have been extended to the motion recognition area. Because the estimation of PPIs through protein complexes is primarily determined by evaluating the distances between residues, this has a high commonality with image recognition. In this study, both DenseNet and ResNet backbone structures are implemented in 3D convolutional networks to resolve protein 3D structure data. The proposed method showcases promising results in predicting PPIs, thereby highlighting the potential of 3D space rendering processing methods in advancing structural biology research.

## 2. Results

### 2.1 SpatialPPI prediction pipeline

SpatialPPI is a pipeline that could predict the possibility of two single-chain proteins interacting with each other. The input of the pipeline is the sequence information of two proteins. The structure information of the potential complex will be predicted by Alphafold Multimer, and then the prediction result will be rendered into 3D tensors as the input to the convolutional neural network to donate the prediction. Figure 1 shows the flow chart of SpatialPPI.

**Figure 1.**
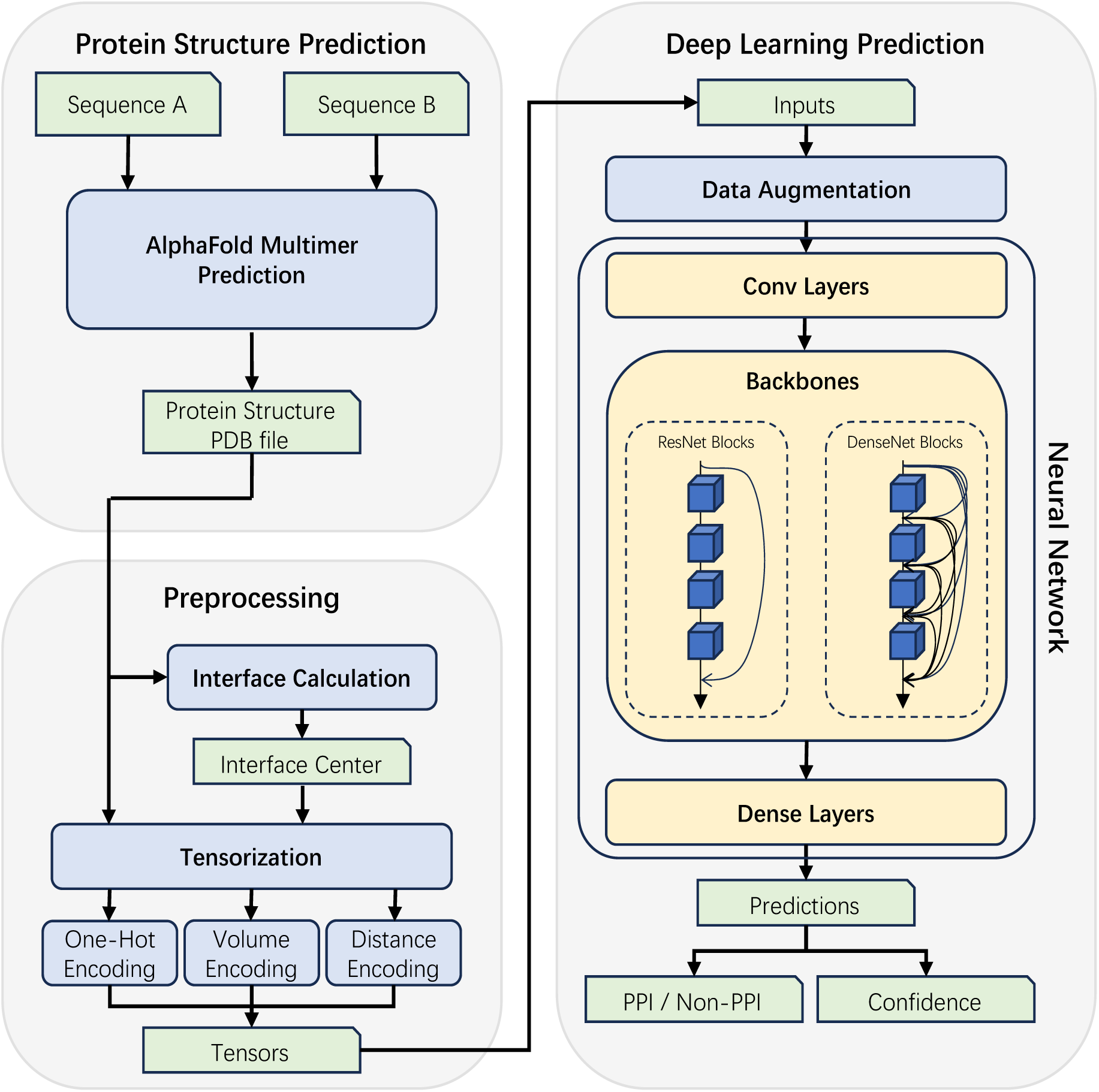
Flow chart of SpatialPPI. Firstly, AlphaFold Multimer predicts the two input protein sequences and generates the structure of the protein complex, which is stored in a PDB file. Tensorize the protein complex by calculating its interface using one of one-hot encoding, volume encoding, or distance encoding. The tensor is then passed to the neural network as input data. After data augmentation, the DenseNet backbone or ResNet backbone predicts and outputs whether the two proteins interact and the confidence of the prediction.

### 2.2 Evaluations between different backbone and tensorization methods

In order to evaluate the performance of the model, multiple sets of tests were conducted. These tests encompassed various experiments involving comparing different backbone architectures and tensorization methods, a comparative analysis against other state-of-the-art PPI prediction methods, and a completely independent supplementary test dataset was also used to validate the results. Regarding the evaluation criteria, binary accuracy (ACC), area under curve (AUC), precision, and recall were calculated for the evaluation. Moreover, the model’s performance was visually represented by generating receiver operating characteristic (ROC) curves.

SpatialPPI performed 5-fold cross-validation on the dataset using four different sets of parameters, where the dataset was partitioned into five mutually exclusive subsets. In each fold, four of these subsets were used as the training set, while the remaining one served as the test set. By aggregating the predictions made on the test set over five iterations, a comprehensive prediction for the dataset was generated. The test results are shown in Table 1 and Figure 2.

**Figure 2.**
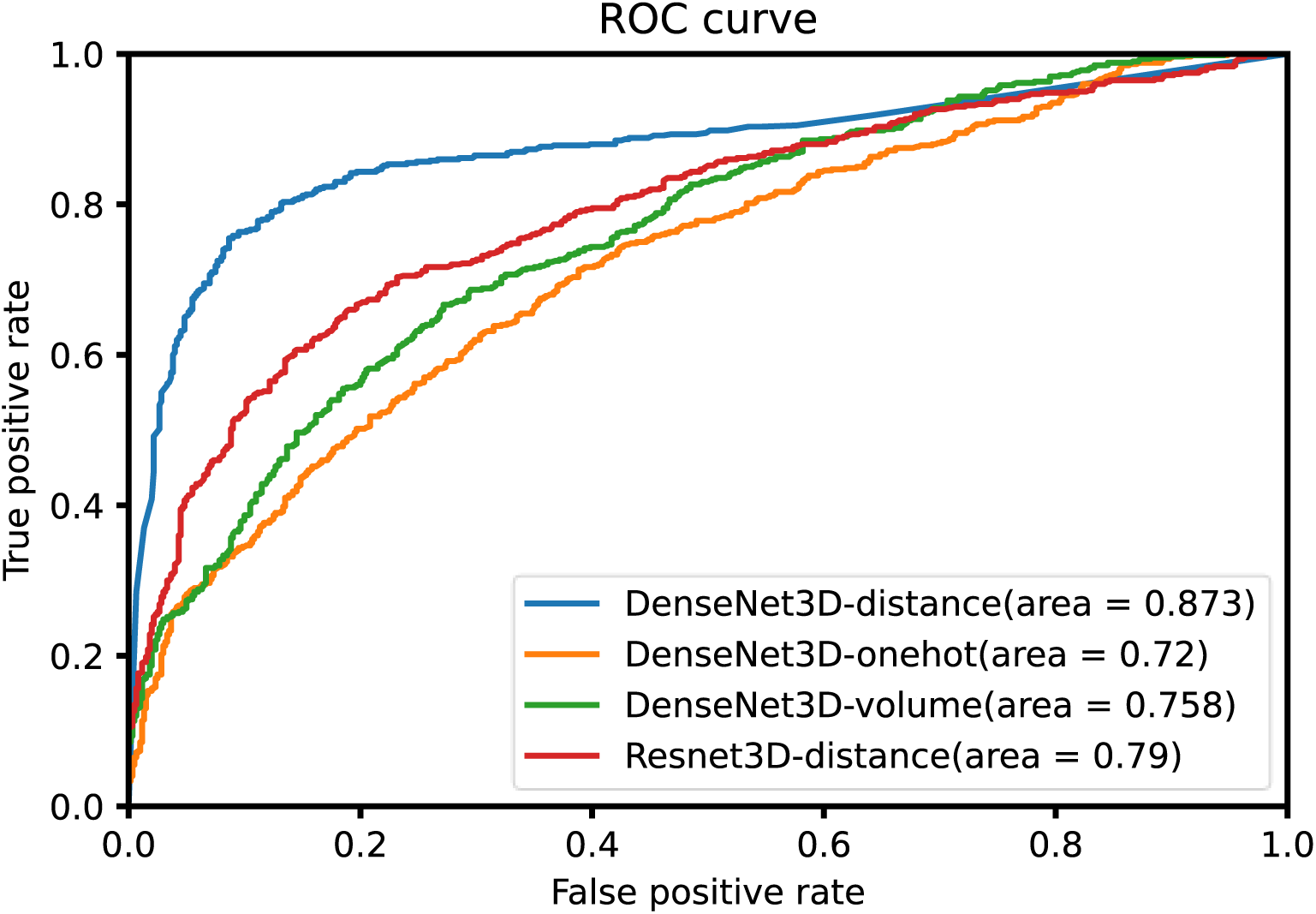
The receiver operating characteristic (ROC) curve for different encoding methods and network backbones based on the 5-fold cross-validation results of the dataset.

**Table 1.** Comparison of accuracy (ACC), area under the receiver operating characteristic curve (AUC), precision, and recall for different encoding methods and network backbones based on the 5-fold cross-validation results of the dataset.

| Network backbone | DenseNet3D | DenseNet3D | DenseNet3D | Resnet3D |
| --- | --- | --- | --- | --- |
| Tensorization | Distance Encoding | One-Hot Encoding | Volume Encoding | Distance Encoding |
| ACC | **0.827** | 0.650 | 0.666 | 0.728 |
| AUC | **0.873** | 0.720 | 0.758 | 0.790 |
| precision | **0.837** | 0.709 | 0.636 | 0.813 |
| recall | **0.813** | 0.508 | 0.773 | 0.593 |

The utilization of DensetNet3D as the backbone architecture along with the distance tensorization method yielded the best performance across all metrics. In contrast, the performance of the one-hot encoding method was less satisfactory. This result confirms that distance encoding provides a practical reference for the neural network’s attention to interatomic distances. It also demonstrates its spatially dense characteristics, which enable more comprehensive updates of neural unit weights. Notably, the model using ResNet3D as the backbone architecture has a precision close to the DensetNet3D model. However, the recall of the ResNet3D model could have been more satisfactory. This implies that the ResNet3D model can effectively identify true negative data with relatively reasonable accuracy but struggles to detect true positive data. In other words, the structure of DenseNet3D allows the network to retain some distance information in the input data during transmission, thereby improving performance. Consequently, the combination of distance encoding and DensetNet3D backbone was selected as the top-performing configuration to represent the performance of SpatialPPI for further analysis and evaluation.

### 2.3 Comparison with the existing PPI prediction method

In this study, four stat-of-art PPI prediction methods have been used as comparison objects to evaluate the performance of the model. Those methods are D-script [31], DeepTrio [32], PEPPI [22] and SpeedPPI [33].

D-script is a deep learning method for directly predicting PPIs from protein sequences. D-script achieves this by utilizing a natural language processing approach to design a pre-trained language model for generating input representations of protein sequences. D-script then estimates the protein’s contact map and uses an interaction module to summarize the interactions. This method is highly user-friendly. Currently, two pre-trained models are available: the Human D-SCRIPT model (from the original D-SCRIPT paper) and the Human Topsy-Turvy model [34] (recommended by developers). Both of these models are evaluated in this comparison.

DeepTrio predicts PPIs by inputting protein sequences into the Siamese architecture with mask multiple parallel convolutional neural networks. It also provides visualization of the importance of each protein residue in both online and offline tools, along with additional predictions for single proteins. The predictive results of DeepTrio are obtained through a 5-fold validation process on the dataset.

PEPPI uses a multi-prong pipeline to predict PPIs by Gaussian kernel density estimation. These modules include a database lookup module, a conjoint triad-trained neural network, and two “interology” based modules: a threading-based module using a modified version of SPRING [35] and a sequence-based module using BLAST [36].

SpeedPPI is a PPI prediction method based on predicted protein structures [37]. According to the introduction by the FoldDock team, SpeedPPI is an improved version of FoldDock [20], with enhancements in usability and speed [37]. This method relies on the pLDDT scores of predicted interaction residues obtained from AlphaFold. By employing a sigmoidal projection, SpeedPPI estimates pDockQ scores, which are used to assess interactions. We used the protein complex structures predicted by AlphaFold Multimer in the SpatialPPI pipeline for SpeedPPI analysis.

Table 2 and Figure 3 show the predicted results for each method and SpatialPPI. Overall, SpatialPPI exhibits outstanding performance compared to other similar methods. SpatialPPI achieves the highest accuracy, area under the curve, and recall among all methods. While PEPPI and original D-script model display high precision, their ability to detect true positives is less satisfactory. It is hypothesized that the training process of the network using conjoint triad features in PEPPI, which shares data sources with the negative dataset in this study, and the inclusion of all proteins in both negative and positive datasets used in this research may have led to an emphasis on protein sequence characteristics over the analysis of protein-protein associations, resulting in such prediction outcomes. In contrast, the updated D-script Human Topsy-Turvy model exhibits a more balanced performance. DeepTrio performs well overall and boasts the fastest computational speed among all methods. Similar to PEPPI and D-script with human weights, it has a relatively lower but acceptable recall. SpeedPPI’s performance relies entirely on AlphaFold’s predictive capability because it is based on the mapping of pLDDT scores. Additionally, SpeedPPI may be influenced to some extent due to not specifically handling the calculation of average pLDDT scores for long single-chain area that are not structurally entangled with another protein.

**Figure 3.**
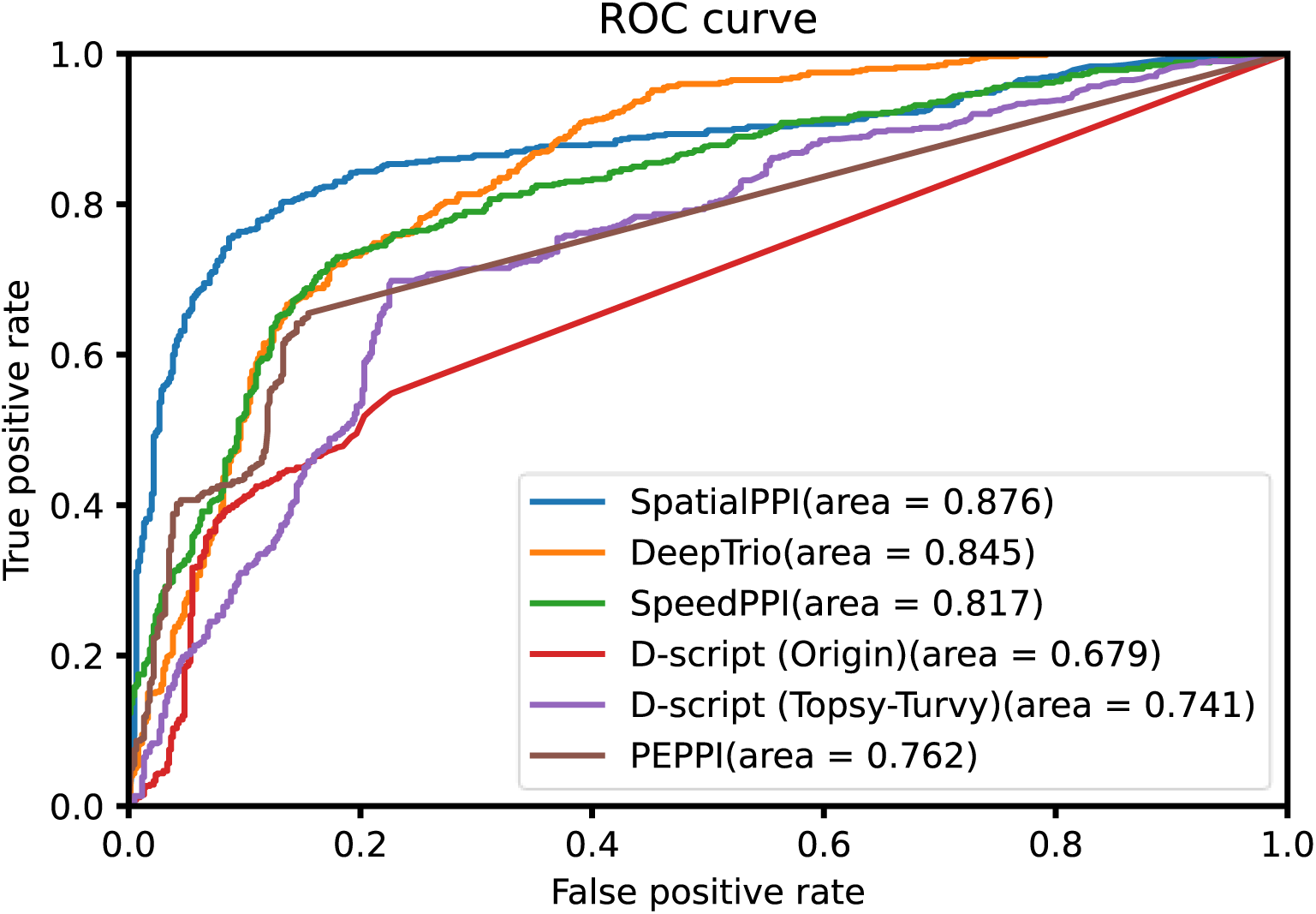
The receiver operating characteristic (ROC) curve for SpatialPPI, DeepTrio, SpeedPPI, PEPPI and D-script.

**Table 2.** Comparison of accuracy (ACC), the receiver operating characteristic curve (AUC), precision, and recall for SpatialPPI, DeepTrio, SpeedPPI, PEPPI and D-script.

|  | SpatialPPI | DeepTrio | SpeedPPI | D-script<br>(Origin) | D-script<br>(Topsy-Turvy) | PEPPI |
| --- | --- | --- | --- | --- | --- | --- |
| ACC | 0.827 | 0.765 | 0.773 | 0.631 | 0.736 | 0.680 |
| AUC | 0.876 | 0.845 | 0.817 | 0.679 | 0.741 | 0.762 |
| precision | 0.837 | 0.830 | 0.807 | 0.852 | 0.755 | 0.906 |
| recall | 0.813 | 0.667 | 0.718 | 0.317 | 0.698 | 0.402 |

### 2.4 Evaluations on additional dataset

To validate the robustness of the network, experiments on an additional dataset was conducted. The additional dataset is a subset of the data utilized in the DeepTrio research [38]. Positive pairs were selected from human species data obtained from BioGRID. Moreover, negative data was generated by shuffling one sequence from the positive pair. Since the particularity of amino acid sequences, the probability that the shuffled proteins can still interact is negligible, and this method was demonstrated by Kandel *et al.*[39]. Specifically, 571 non-repeating positive pairs and their corresponding shuffled counterparts are chosen as the negative dataset. For the execution of SpatialPPI, DenseNet3D backbone and distance encoding were used to perform 5-fold validation on the additional dataset. Furthermore, D-script (Topsy-Turvy), DeepTrio, and SpeedPPI were also tested on the additional dataset.

The prediction results were shown in Table 3. The comparison results indicate that SpatialPPI outperforms other methods, and demonstrates that SpatialPPI’s ability to learn the spatial structural relationships of residues is also applicable to other data sets. Additionally, a significant difference was observed in performance on the negative dataset compared to the dataset based on Negatome 2.0. Almost all methods showed a notable improvement in accuracy on the negative dataset portion. This may suggest that the manner in which the negative dataset was generated, while theoretically non-interacting, exhibits substantial differences in sequence characteristics from naturally occurring proteins. Consequently, these differences make the negative dataset more easily distinguishable in practice.

**Table 3.**
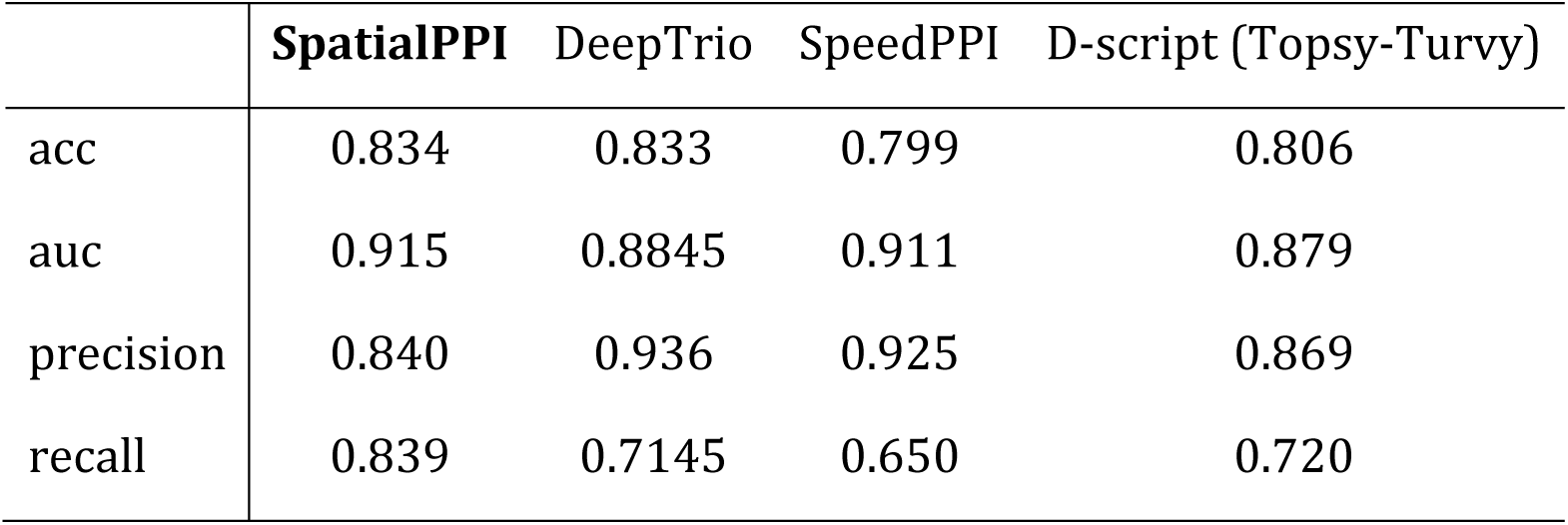
Comparison of accuracy(ACC), area under curve (AUC), precision, and recall for SpatialPPI, DeepTrio, SpeedPPI and D-script using the Topsy-Turvy model, performed on the additional dataset from DeepTrio’s research.

## 3. Discussion and Conclusions

To be noticed that due to current limitations in GPU computational power, the execution speed of this pipeline could be faster. As a result, the amount of data used in the experiments is smaller than in other deep learning studies. Additionally, this implies that the network’s composition is primarily influenced by the protein interfaces that are already present in the dataset. However, the model’s performance on the supplementary training dataset demonstrates its reasonable robustness. The model is believed to have reasonable performance on other datasets with fine-tuning.

Although both negative PPIs (obtained from Negatome 2.0 database) and positive PPIs (obtained from BioGRID database) are experimentally determined data, it should be noted that conflicts exist between them, i.e. some PPIs are recorded in both. This may be due to experimental errors or incomplete statistics. Negatome 2.0 is one of the most widely used negative PPI database, and it has been 9 years since its release. During this period, as technology improved and experiments were carried out, the understanding of PPIs has become more profound, and some of the data may have changed. For current PPI studies using computational methods, these ambiguous data should be removed to ensure the reliability of the dataset.

By calculating the proportion of valid values in the input data, we define the fill rate of the input neural network data as the number of non-zero values in the input tensor divided by the input size. For the three tensorization methods used in this study, their average fill rates in the central area are: one-hot encoding at 3%, volume encoding at 20%, and distance encoding at 95%. In other studies, there are also applications of adapting Gaussian blur in one-hot encoding. That method has a similar fill rate of volume encoding, but lacks spatial distribution information of atoms compared with volume encoding. The distance-based encoding method could fill the spatial sparsity in protein structures, enabling a more comprehensive update of neurons in convolutional neural networks, thereby improving the model performance.

Although AlphaFold Multimer achieves unprecedented high accuracy in predicting protein complex structures, it could still output prediction results with high pLDDT scores even when the input protein sequences cannot interact with each other. By exploiting this feature, we were able to generate a three-dimensional spatial model of the negative protein data set, thereby providing protein structural data for neural network analysis of PPIs. The benefit is that SpatialPPI can use protein structural information to predict PPIs even for proteins whose three-dimensional structure has not been experimentally determined. On the other hand, since intrinsically disordered regions in proteins are represented by a ribbon-like appearance, which should be interpreted as a prediction of a disordered state and not as structural information, their spatial contact with another protein should be ignored. By introducing a process that removes disordered areas, SpatialPPI can correctly distinguish PPIs without being interfered with by entangled disordered areas, which is critical to SpatialPPI’s classification capabilities.

In conclusion, SpatialPPI is a pipeline for predicting PPIs by utilizing protein complex structures predicted using AlphaFold Multimer. Through testing and evaluating three strategies for rendering protein structure information into spatial tensors and two commonly used backbones for image recognition and video classification, this research provides a reference for analyzing protein three-dimensional structures using neural networks. Moreover, this underscores the fact that both protein structure data and image data share a spatially distributed information basis, which results in their similarity in terms of features and allows them to be captured and classified by similar convolutional neural network architectures. Furthermore, the necessity of using experimentally validated negative datasets is demonstrated through testing on two independent datasets. With the advancement in computational capabilities, it is hoped that this analytical approach can further develop and contribute to the understanding of the principles underlying PPIs.

## 4. Materials and Method

### 4.1 Dataset Construction

The quality and reliability of PPI data are of paramount importance for the development and evaluation of computational models and algorithms for PPI prediction. In this study, we used BioGRID[40], a comprehensive biomedical interaction repository, as the primary source of interacting protein pairs. BioGRID contains over 2 million experimentally validated PPIs from more than 70,000 publications in the primary literature. This ensures the authenticity and accuracy of the positive dataset used in this study.

In contrast, obtaining reliable negative datasets for PPI prediction remains a challenging task. As demonstrated by Wei *et al.* [41], experimentally validated datasets of non-interacting proteins are more reliable as negative datasets than randomly matching proteins that are not present in the positive set. To address this issue, we employed Negatome 2.0[42], a manually annotated literature data resource, to source the non-interacting protein pairs for our study. Negatome 2.0 is a dataset that contains protein-protein pairs which are unlikely to engage in physical interactions, and includes 2171 non-interacting pairs from 1828 proteins in the manual subset which is manually annotated data collected from literatures.

However, it should be noted that conflicts exist between the positive and negative datasets. There are 26.5% of pairs from Negatome have conflicting records in BioGRID, of which 296 conflicts have more than one publication in BioGRID. To minimize the impact of such conflicts on the quality of the resultant dataset and improve the quality of the dataset, we meticulously curated the dataset by removing conflicts, duplicates, and self-interactions. Sequences with high identities were also removed. This is to prevent when sequences with high similarity appear in both the training set and the test set, the evaluation results will be affected by over-fitting and fail to reflect the true performance of the model.

Moreover, we ensured a balanced and diverse representation of the dataset by requiring each protein to have at least one protein pair in which two proteins interact with each other and one protein pair in which two proteins do not interact with each other.

The final dataset comprises a total of 1200 protein pairs, consisting of 600 positive and 600 negative instances, derived from 375 Homo sapiens proteins. Overall, the resultant dataset represents a high-quality and reliable resource for the development and evaluation of computational models and algorithms for PPI prediction.

### 4.2 Protein Structure Using AlphaFold Multimer

In view of the achievement of AlphaFold [43] in the 14th Critical Assessment of Structural Prediction competition [44], the algorithm was used in this study for generating possible protein complexes of the input sequences. AlphaFold Multimer is a deep neural network-based method developed by the AlphaFold team at DeepMind to predict the structures of protein complexes, which are composed of multiple interacting proteins. The method uses the AlphaFold model to predict the structures of individual protein subunits and then assembles them into a complex using an optimization algorithm. The optimization algorithm considers both the predicted structure and the predicted binding affinity of each subunit to minimize the energy of the complex. This high-accuracy protein structure prediction method can make the pipeline only need to accept easy-to-acquire protein sequence data, and at the same time obtain more spatial information through the protein structure to improve prediction accuracy.

Structural prediction of 1200 protein complexes was performed by using AlphaFold Multimer version 2.3.1 released on Jan 12, 2023 [45]. AlphaFold multimer was executed with using the jackhmmer search from HMMER3[46] targeting Uniref90 [47], UniProt [15], and MGnify [48] and a joint search on the Big Fantastic Database [49, 50] and UniRef30 [47] using HHBlits [51]. For each input, five different models was generated by AlphaFold Multimer, and each model donated one prediction, yielding a total of 6000 PDB files.

### 4.3 3D Rendering of Protein Structure

Extracting the features of input data and converting them into data structures that can be recognized by neural networks has always been an issue that has attracted much attention in deep learning research. The prediction results of AlphaFold Multimer are in PDB format [52], which is a list of coordinates for atoms. The method used in this study is to map the preprocessed protein complex structure into a three-dimensional tensor according to the coordinates. This method has also been used in studies such as [24, 53]. Its advantage is that compared with linear data stored in PDB format files, or two-dimensional data such as distance matrices, it can better express the spatial structure information of proteins, thus providing more information for the input part of the neural network. And have the possibility to conclude details that may not have been summarized by human analysis. The current opinion is that whether proteins can bind to each other mainly depends on the analysis of the contact surface of the two proteins [54]. Therefore, in this study, the contact surface part of the protein complex was selected as the center of the input data. To this end, the preprocessing stage consists of removing the predicted low-confidence parts, calculating the residues belonging to the contact surface, and then building a tensor centered on the geometric center of the contact surface.

Disordered structures are common in eukaryotic proteomes. Previous work estimated the proportion of disordered residues in the human proteome at 37-50% [56]. Examples of disordered areas are shown in Figure 4. The AlphaFold research team found that pLDDT is an adequate predictor of intrinsically disordered area, because the distribution of pLDDT between resolved and unresolved residues in the PDB sequence is very different [57]. Currently, AlphaFold puts long domains with pLDDT scores less than 50 to exhibit a ribbon-like appearance and should be interpreted as a prediction of a disordered state and not as structural information. Hence, these parts have no substantial effect on the structure of PPIs. However, these free, long, single chains are often entangled with another chain to form a part of the region very close to each other. Therefore, long single strands with pLDDT scores less than the threshold 50 will be deleted to reduce the influence of contact regions other than the interface predicted by AlphaFold.

**Figure 4.**
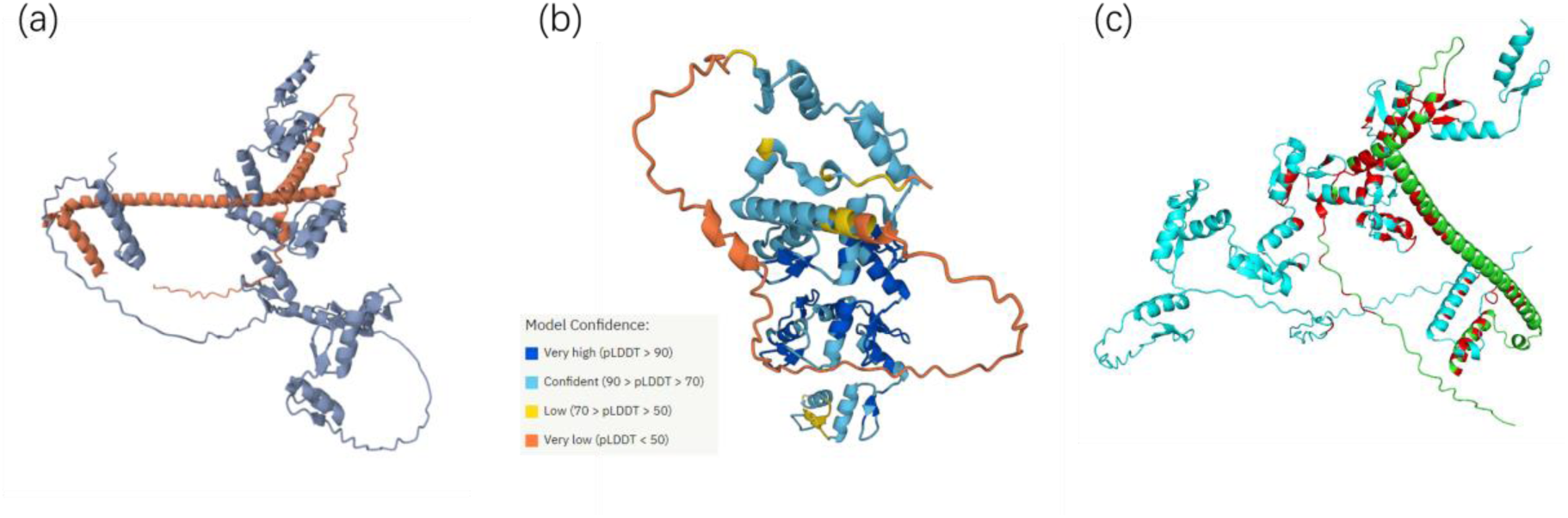
Examples of disordered areas, which are usually appear in the Alphafold prediction result. (a) AlphaFold Multimer predicted structure of protein complex of P18847 and Q9NQX6, (b) Predicted pLDDT score of Q9NQX6[55], (c) Contact area of (a) with the threshold of 8 Å, marked as red. In addition, this protein pair is labeled as negative from Negatome 2.0.

The center of the tensor is defined as the geometric center of the contacted residue from two chains. The interface calculation is mainly by calculating the distance matrix of carbon atoms with remoteness indicator code alpha from the PDB file. The threshold for the two protein-contact regions was set at 12 Å rather than 8 Å, which is commonly used [58]. Considering the error range in AlphaFold’s prediction, more possible regions are expected to be selected for reference. If all distances in both sequences are greater than the threshold, the threshold is gradually increased until at least one residue is selected for each of the two sequences. The reason is to choose an area where the two sequences are spatially closest to each other as the center of the input data, thus ensuring that the input data provides the most possibility for subsequent neural network analysis.

Each tensor is defined to represent a three-dimensional space, and each position in the matrix represents the spatial information in a cube with a side length of 1 Å. The tensor size has been restricted to [64, 64, 64, 8]. Due to the limitations of computing power in the current era, it is practically impossible to continue to reduce the per-unit length since it will cost dozens of times. The first three indexes are coordinates rounded to 1 Å, which means each tensor stands for a space of cube with size 64 Å × 64 Å × 64 Å. The last dimension is for four kinds of atoms: carbon, nitrogen, oxygen, and sulfur. The four elements located in the two amino acid sequences occupy 4 positions respectively, totaling 8 positions. The hydrogen was excluded since they are not the main structure predicted by AlphaFold but are filled in as supplementary.

After defining the tensor, the next step is to calculate the representation of each atom in the tensor. Three kinds of tensorization methods were implemented and evaluated: one-hot encoding, volume encoding, and distance encoding.

#### 4.3.1 One-hot Encoding

The most commonly used encoding method is one-hot encoding. This method begins by rounding the coordinates of an atom to an integer to determine where the atom is located. Then, the value of that cell is marked as 1 according to the chain and type of the atom, and all other positions are marked as 0. This method was used in the study by Jones *et al*. [24] Schematic diagram of One-hot encoding shows in Figure 5(a).

**Figure 5.**
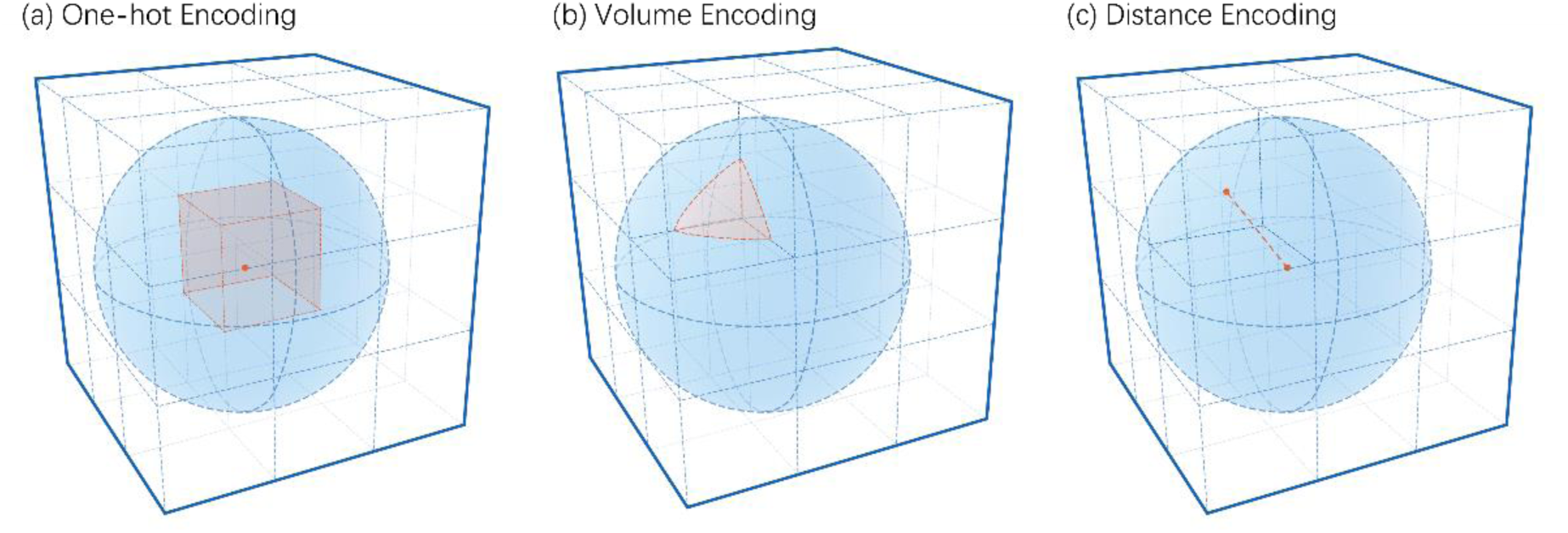
Schematic diagram of three encoding methods for rendering atoms into space. (a) One-hot Encoding: Mark the cell where the atom center is located, (b) Volume Encoding: Values of each cell equal to the volume of the intersection of the cell and the atom, (c) Distance Encoding: Values of each cell equal to the distance from the atom center to the cell.

The advantage of this method is that it is easy to calculate and understand. The spatial distribution of atoms is expressed in the most direct way. Nevertheless, there is also the problem of deleting all the atomic coordinate information after the decimal place. Consider that it is generally believed that the condition for the contact of two residues is that their ca atomic distance is less than 8 Å, but the discarded part may be closer to 1 Å. Such a high deviation may have a harmful effect on the forecast results.

#### 4.3.2 Volume Encoding

Volume encoding is a more accurate way to express the spatial distribution of atoms than one-hot encoding since the atomic radius may not be ignorable. A table of the diameter of the atoms contained in protein shows in Table 4. It calculates the volume distribution of atoms and defines the value in each cell as the intersection of the atomic volume and the cell. Schematic diagram of One-hot encoding shows in Figure 5(b). This method could be more accurate in spatial representation than one-hot. Because the probability that the center of an atom happens to be in the center of a cell is negligible, this method provides a more explicit representation of the spatial distribution of atoms without increasing the input size of the neural network.

**Table 4.** The diameter of the atoms included in PDB files[59]. The atomic diameters of the main atoms constituting amino acids: carbon, nitrogen, oxygen and sulfur are all larger than the designed unit side length, which is 1 Å.

| Atom | Atomic Diameter (Å) |
| --- | --- |
| Hydrogen | 0.50 |
| Carbon | 1.40 |
| Nitrogen | 1.30 |
| Oxygen | 1.20 |
| Sulfur | 2.00 |

For a cell at coordinate (*x*_*i*_, *y*_*j*_, *z*_*k*_), and side length is equal to *a*, and the center of the atom is defined as (*x*_*a*_, *y*_*a*_, *z*_*a*_) with radius *R*. Let *S* represent the projection of the intersection between the cell and the atom onto the *x* − *y* plane. Then the volume of the intersection could be calculated by the following formula:

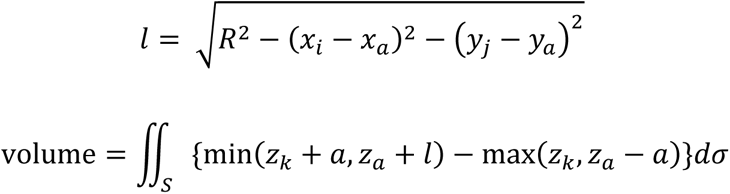

The implementation of the calculation of the volume for the intersection of the atom and the unit would be calculated by integration. However, it is very time-consuming. Therefore, a random simulation is used to replace it. The random simulation method generates 1,000,000 random points inside the cell. By counting the number of points inside the atom, the volume of the intersection could be estimated. The volume calculated in this method is compared with the volume calculated by integration to verify its effectiveness, and the accuracy of the method is about 99.5%.

#### 4.3.3 Distance Encoding

Distance encoding is based on the characteristics of interatomic interaction. The main factors that affect PPIs include electrostatic interaction, hydrogen bonds, hydrophobic effect, and van der Waals forces [60]. The main factor affecting these factors is distance.

The distance encoding algorithm renders the surrounding cells as the distance between that cell and the center of the atom, starting from each atomic position. One problem with both the one-hot encoding and volume encoding is that most of the space is empty when represented in space due to the nature of the protein complex itself. This property causes the resulting tensor to be mathematically treated as a sparse matrix. Studies have shown that the performance of neural networks deteriorates when the input matrix is too sparse [61]. This is because such a sparse matrix results in each input data providing updates to only a limited number of neurons, then dramatically reducing training efficiency. At the same time, this sparse matrix will cause the trained network to rely more on a few local details, thus reducing the stability of the network. The advantage of distance coding is that it fills most of the space with data, making tensor spatially dense. This method provides relatively wealthy atomic coordinate information without increasing the amount of data as volume encoding and improves the training efficiency and quality of the neural network.

The calculation of distance is defined as the Euclidean Distance from the center of the cell to the center point of the atom. The maximum distance for rendering is set at 12 Å, which is the same as the interface calculation. When a cell is within the threshold of two or more identical atoms at the same time, the value of the cell is defined as the smaller value of the two distances. When a cell is within the threshold of two or more different atoms at the same time, the distance between the corresponding atoms is marked at the same time. For a cell at coordinate (*x*_*i*_, *y*_*j*_, *z*_*k*_), and the center of the atom is defined as (*x*_*a*_, *y*_*a*_, *z*_*a*_), then the distance is defined by the following formula:

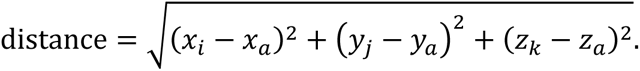

### 4.4 Architecture of Neural Networks

Convolutional neural networks have been widely used in computer vision [25–28]. Specifically, the Deep Residual Network (ResNet)[30] and the Densely Connected Convolutional Networks (DenseNet)[29] have become the mainstream backbones in computer vision networks [62, 63]. Both of these network structures are designed to solve the degradation problem that occurs when the depth of the network structure increases, that is, the accuracy of the network saturates or even decreases as the depth of the network increases. Protein structure analysis shares significant similarities with computer vision regarding data structure. The image information is represented as a three-dimensional array in computing: the first two dimensions represent its spatial coordinates, and the third dimension represents the color information of position. Furthermore, the techniques used in video analysis are even much closer to protein processing. By stacking multiple images on top of each other, the first dimension represents the time, the second and third dimensions represent the spatial coordinates, and the last dimension is the color information at that position. In deep learning, the core issue of computer vision analysis and protein structure analysis is similar: finding regions with features by calculating correlations between adjacent units in space, detecting patterns in these regions, and classifying them based on their features. Due to their high similarity, it is expected that ResNet and DenseNet can also leverage their advantages in protein analysis [64, 65].

The number of samples affects the model performance. However, due to the specificity of biological data, many data augmentation techniques are inappropriate for this study, such as flipping [66]. Therefore, data augmentation relies on rotations of the input data. Since the input tensor is defined as a cube, the coordinate system could be established from eight vertices of the cube as coordinate origins, and interchanging the *x*, *y*, and *z* axes results in a total of 24 rotation possibilities. This approach alleviates the data shortage. At the same time, the network structure is trained in balance by rotation to deal with the inconsistent orientation of the predicted protein complex.

Modified ResNet and DenseNet were used as options of backbones for the neural network in this study, with implementations adapted from Ju’s work [67] and Dudovitch’s work [68]. Both ResNet and DenseNet are composed of multiple blocks with similar structures. Each block includes convolutional layers, activation layers, pooling layers, and dropout layers. The convolutional layers within the same block have the same size, and different blocks are connected by transition blocks to reduce the size. ResNet introduces shortcut connections in each block, passing data from the front of a block directly to the back to mitigate the gradient vanishing problem, thereby improving the performance of deep neural networks. DenseNet follows a similar idea to ResNet but connects all previous layers to subsequent layers. Figure 6 shows the diagram of the network architecture. Finally, the output of the entire network is a 2-length array, representing the probabilities of the input data being positive and negative, respectively. The higher of the two probabilities is defined as the network’s prediction, while the difference between these two probabilities is defined as the model’s confidence in the prediction. For the loss function, positive data is labeled as [0, 1], negative data as [1, 0], and the weights are updated by calculating the categorical cross-entropy between the predicted results and the labels.

**Figure 6.**
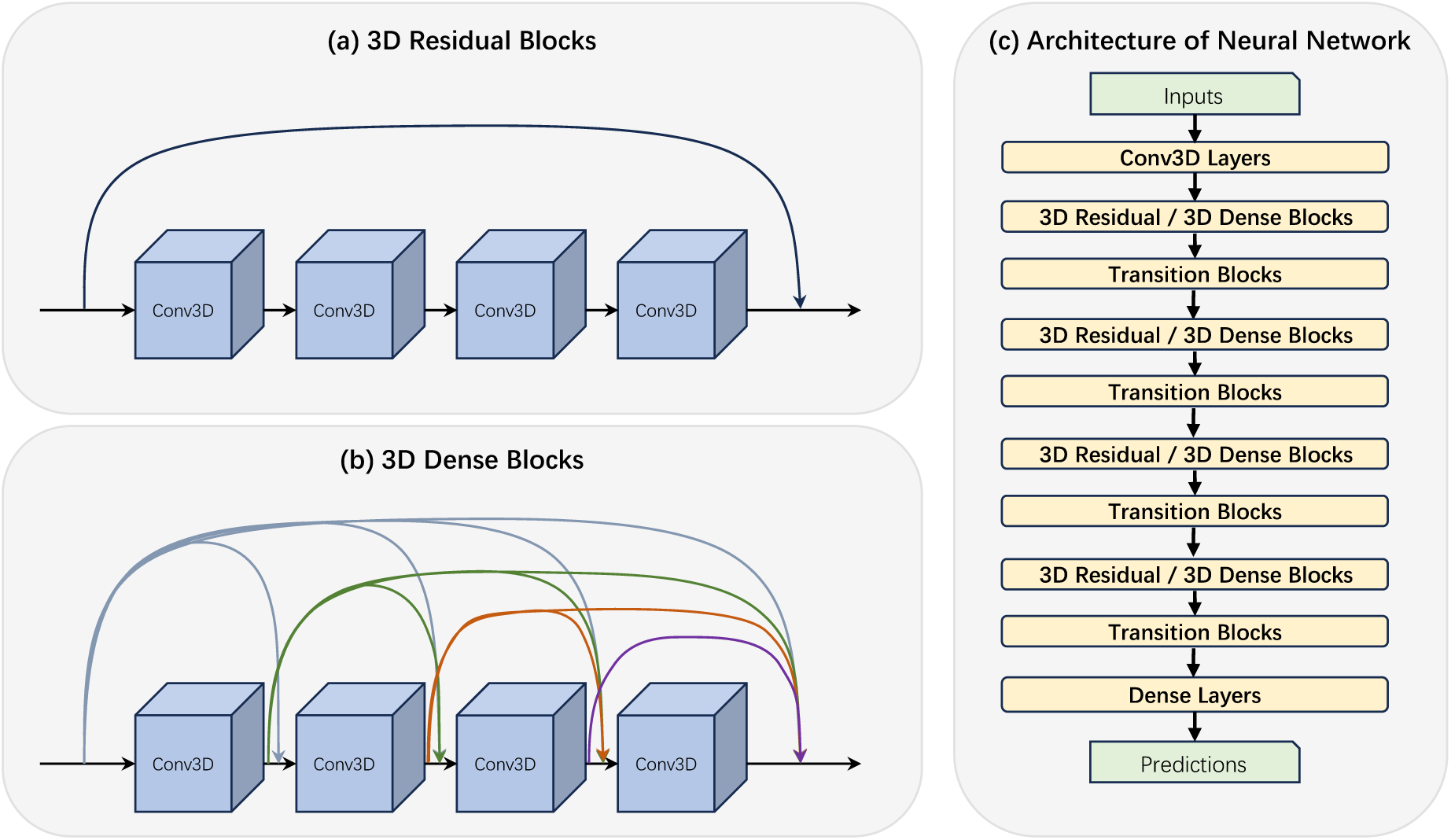
Details of the structure of the SpatialPPI network. (a) Structure of 3D Residual Blocks. In each residual block, the input data is directly accumulated into the output layer through a shortcut. (b) Structure of 3D Dense Blocks. For dense blocks, the input data and the output data of each layer of the convolutional network except the last layer are passed backward using shortcut. (c) Architecture of the Neural Network. Multiple backbone blocks are gradually reduced in size after being connected using transition blocks, and finally dense layers conclude and output the prediction.

## Competing interests

All authors declare no financial or non-financial competing interests.

## Acknowledgements

M.O. discloses support for the research of this work from JST FOREST [grant number JPMJFR216J], and AMED BINDS [grant number JP23ama121026].

## Code availability

The underlying code and datasets for this study is available in GitHub and can be accessed via this link https://github.com/ohuelab/SpatialPPI.

## Contributions

W.H. and M.O. conceived the study; W.H. designed code; W.H. performed calculations; W.H. and M.O. analyzed data; W.H. wrote the initial manuscript; All authors approved the final version of the manuscript.

## Notes

### Competing Interest Statement

The authors have declared no competing interest.

## Reference

1. Dunham, B., & Ganapathiraju, M. K. (2021). Benchmark evaluation of protein–protein interaction prediction algorithms. Molecules, 27(1), 41. doi10.3390/molecules27010041

2. Tsukiyama, S., Hasan, M. M., Fujii, S., & Kurata, H. (2021). LSTM-PHV: Prediction of human-virus protein–protein interactions by LSTM with word2vec. Briefings in Bioinformatics, 22(6). doi10.1093/bib/bbab228

3. Sun, T., Zhou, B., Lai, L., & Pei, J. (2017). Sequence-based prediction of protein protein interaction using a deep-learning algorithm. BMC Bioinformatics, 18(1). doi10.1186/s12859-017-1700-2

4. Murakami, Y., & Mizuguchi, K. (2017). PSOPIA: Toward more reliable protein-protein interaction prediction from sequence information. 2017 International Conference on Intelligent Informatics and Biomedical Sciences (ICIIBMS). doi10.1109/iciibms.2017.8279749

5. Baspinar, A., Cukuroglu, E., Nussinov, R., Keskin, O., & Gursoy, A. (2014). Prism: A web server and repository for prediction of protein–protein interactions and modeling their 3D complexes. Nucleic Acids Research, 42(W1). doi10.1093/nar/gku397

6. Baranwal, M., Magner, A., Saldinger, J., Turali-Emre, E. S., Elvati, P., Kozarekar, S., VanEpps, J. S., Kotov, N. A., Violi, A., & Hero, A. O. (2022). Struct2Graph: A graph attention network for structure based predictions of protein–protein interactions. BMC Bioinformatics, 23(1). doi10.1186/s12859-022-04910-9

7. Murakami, Y., & Mizuguchi, K. (2014). Homology-based prediction of interactions between proteins using averaged one-dependence estimators. BMC Bioinformatics, 15(1). doi10.1186/1471-2105-15-213

8. Chen, C.-C., Lin, C.-Y., Lo, Y.-S., & Yang, J.-M. (2009). PPISearch: A web server for searching homologous protein–protein interactions across multiple species. Nucleic Acids Research, 37(suppl_2). doi10.1093/nar/gkp309

9. Hashemifar, S., Neyshabur, B., Khan, A. A., & Xu, J. (2018). Predicting protein–protein interactions through sequence-based deep learning. Bioinformatics, 34(17), i802–i810. doi10.1093/bioinformatics/bty573

10. Yao, Y., Du, X., Diao, Y., & Zhu, H. (2019). An integration of deep learning with feature embedding for protein–protein interaction prediction. PeerJ, 7. doi10.7717/peerj.7126

11. Chen, M., Ju, C. J., Zhou, G., Chen, X., Zhang, T., Chang, K.-W., Zaniolo, C., & Wang, W. (2019). Multifaceted protein–protein interaction prediction based on Siamese residual RCNN. Bioinformatics, 35(14), i305–i314. doi10.1093/bioinformatics/btz328

12. Li, H., Gong, X.-J., Yu, H., & Zhou, C. (2018). Deep neural network based predictions of protein interactions using primary sequences. Molecules, 23(8), 1923. doi10.3390/molecules23081923

13. Schwede, T. (2013). Protein modeling: What happened to the “Protein structure gap”? Structure, 21(9), 1531–1540. doi10.1016/j.str.2013.08.007

14. Berman, H., Henrick, K., & Nakamura, H. (2003). Announcing the worldwide Protein Data Bank. Nature Structural & Molecular Biology, 10(12), 980–980. doi10.1038/nsb1203-980

15. Bateman, A., Martin, M.-J., Orchard, S., Magrane, M., Ahmad, S., Alpi, E., Bowler-Barnett, E. H., Britto, R., Bye-A-Jee, H., Cukura, A., Denny, P., Dogan, T., Ebenezer, T., Fan, J., Garmiri, P., da Costa Gonzales, L. J., Hatton-Ellis, E., Hussein, A., Ignatchenko, A., … Zhang, J. (2022). Uniprot: The Universal Protein Knowledgebase in 2023. Nucleic Acids Research, 51(D1). doi10.1093/nar/gkac1052

16. Soleymani, F., Paquet, E., Viktor, H., Michalowski, W., & Spinello, D. (2022a). Protein–protein interaction prediction with Deep Learning: A Comprehensive Review. Computational and Structural Biotechnology Journal, 20, 5316–5341. doi10.1016/j.csbj.2022.08.070

17. Kundrotas, P. J., Zhu, Z., Janin, J., & Vakser, I. A. (2012). Templates are available to model nearly all complexes of structurally characterized proteins. Proceedings of the National Academy of Sciences, 109(24), 9438–9441. doi10.1073/pnas.1200678109

18. Shoemaker, B. A., & Panchenko, A. R. (2007). Deciphering protein–protein interactions. part II. computational methods to predict protein and domain interaction partners. PLoS Computational Biology, 3(4). doi10.1371/journal.pcbi.0030043

19. Evans, R., O’Neill, M., Pritzel, A., Antropova, N., Senior, A., Green, T., Žídek, A., Bates, R., Blackwell, S., Yim, J., Ronneberger, O., Bodenstein, S., Zielinski, M., Bridgland, A., Potapenko, A., Cowie, A., Tunyasuvunakool, K., Jain, R., Clancy, E., … Hassabis, D. (2021). Protein Complex Prediction with Alphafold-Multimer. doi10.1101/2021.10.04.463034

20. Bryant, P., Pozzati, G., & Elofsson, A. (2022). Improved prediction of protein-protein interactions using AlphaFold2. Nature Communications, 13(1). doi10.1038/s41467-022-28865-w

21. Jumper, J., Evans, R., Pritzel, A., Green, T., Figurnov, M., Ronneberger, O., Tunyasuvunakool, K., Bates, R., Žídek, A., Potapenko, A., Bridgland, A., Meyer, C., Kohl, S. A., Ballard, A. J., Cowie, A., Romera-Paredes, B., Nikolov, S., Jain, R., Adler, J., … Hassabis, D. (2021). Highly accurate protein structure prediction with alphafold. Nature, 596(7873), 583–589. doi10.1038/s41586-021-03819-2

22. Bell, E. W., Schwartz, J. H., Freddolino, P. L., & Zhang, Y. (2022). PEPPI: Whole-proteome protein-protein interaction prediction through structure and sequence similarity, Functional Association, and Machine Learning. Journal of Molecular Biology, 434(11), 167530. doi10.1016/j.jmb.2022.167530

23. Pedregosa, F., Varoquaux, G., Gramfort, A., Michel, V., Thirion, B., Grisel, O., Blondel, M., Prettenhofer, P., Weiss, R., Dubourg, V., Vanderplas, J., Passos, A., Cournapeau, D., Brucher, M., Perrot, M., & Duchesnay, E . Scikit-Learn: Machine learning in Python. Journal of Machine Learning Research.

24. Jones, D., Kim, H., Zhang, X., Zemla, A., Stevenson, G., Bennett, W. F., Kirshner, D., Wong, S. E., Lightstone, F. C., & Allen, J. E. (2021). Improved protein–ligand binding affinity prediction with structure-based deep fusion inference. Journal of Chemical Information and Modeling, 61(4), 1583–1592. doi10.1021/acs.jcim.0c01306

25. Perslev, M., Dam, E. B., Pai, A., & Igel, C. (2019). One network to segment them all: A general, lightweight system for accurate 3D medical image segmentation. Lecture Notes in Computer Science, 30–38. doi10.1007/978-3-030-32245-8_4

26. Nikolaos, A. (2019). Deep learning in medical image analysis: a comparative analysis of multi-modal brain-MRI segmentation with 3D deep neural networks. GitHub. github.com/black0017/MedicalZooPytorch

27. Tran, D., Wang, H., Torresani, L., Ray, J., LeCun, Y., & Paluri, M. (2018, April 12). A closer look at spatiotemporal convolutions for action recognition. CVPR2018

28. Kataoka, H., Wakamiya, T., Hara, K., & Satoh, Y. (2020, April 10). Would mega-scale datasets further enhance spatiotemporal 3D CNNs?. arXiv.org. doi10.48550/arXiv.2004.04968

29. Huang, G., Liu, Z., van der Maaten, L., & Weinberger, K. Q. (2018, January 28). Densely connected Convolutional Networks. CVPR2017

30. He, K., Zhang, X., Ren, S., & Sun, J. (2015, December 10). Deep residual learning for image recognition. CVPR2016

31. Sledzieski, S., Singh, R., Cowen, L., & Berger, B. (2021). D-script translates genome to phenome with sequence-based, structure-aware, genome-scale predictions of protein-protein interactions. Cell Systems, 12(10). doi10.1016/j.cels.2021.08.010

32. Hu, X., Feng, C., Zhou, Y., Harrison, A., & Chen, M. (2021). DeepTrio: A ternary prediction system for protein–protein interaction using mask multiple parallel convolutional neural networks. Bioinformatics, 38(3), 694–702. doi10.1093/bioinformatics/btab737

33. Bryant, P., & Noe, F. (2023). Rapid Protein-Protein Interaction Network Creation from Multiple Sequence Alignments with Deep Learning. doi10.1101/2023.04.15.536993

34. Singh, R., Devkota, K., Sledzieski, S., Berger, B., & Cowen, L. (2022). Topsy-Turvy: Integrating a global view into sequence-based PPI prediction. Bioinformatics, 38(Supplement_1), i264–i272. doi10.1093/bioinformatics/btac258

35. Guerler, A., Govindarajoo, B., & Zhang, Y. (2013). Mapping monomeric threading to protein– protein structure prediction. Journal of Chemical Information and Modeling, 53(3), 717–725. doi10.1021/ci300579r

36. Altschul, S. F., Gish, W., Miller, W., Myers, E. W., & Lipman, D. J. (1990). Basic local alignment search tool. Journal of Molecular Biology, 215(3), 403–410. doi10.1016/s0022-2836(05)80360-2

37. ElofssonLab / FoldDock · GITLAB. GitLab.gitlab.com/ElofssonLab/FoldDock

38. Xiaoti, H. *Huxiaoti/deeptrio*. GitHub. github.com/huxiaoti/deeptrio/tree/master

39. Kandel, D., Matias, Y., Unger, R., & Winkler, P. (1996). Shuffling biological sequences. Discrete Applied Mathematics, 71(1–3), 171–185. doi10.1016/s0166-218x(97)81456-4

40. Stark, C. (2006). BioGRID: A general repository for interaction datasets. Nucleic Acids Research, 34(90001). doi10.1093/nar/gkj109

41. Wei, L., Xing, P., Zeng, J., Chen, J., Su, R., & Guo, F. (2017). Improved prediction of protein– protein interactions using novel negative samples, features, and an ensemble classifier. Artificial Intelligence in Medicine, 83, 67–74. doi10.1016/j.artmed.2017.03.001

42. Blohm, P., Frishman, G., Smialowski, P., Goebels, F., Wachinger, B., Ruepp, A., & Frishman, D. (2013). Negatome 2.0: A database of non-interacting proteins derived by literature mining, manual annotation and protein structure analysis. Nucleic Acids Research, 42(D1). doi10.1093/nar/gkt1079

43. Senior, A. W., Evans, R., Jumper, J., Kirkpatrick, J., Sifre, L., Green, T., Qin, C., Z í dek, A., Nelson, A. W., Bridgland, A., Penedones, H., Petersen, S., Simonyan, K., Crossan, S., Kohli, P., Jones, D. T., Silver, D., Kavukcuoglu, K., & Hassabis, D. (2020). Improved protein structure prediction using potentials from deep learning. Nature, 577(7792), 706–710. doi10.1038/s41586-019-1923-7

44. 14th Community Wide Experiment on the Critical Assessment of Techniques for Protein Structure Prediction. predictioncenter.org/casp14/index.cgi

45. Deepmind. Release Alphafold v2.3.1 · Deepmind/Alphafold. GitHub. github.com/deepmind/alphafold/releases/tag/v2.3.1

46. HMMER. http://hmmer.org/

47. Suzek, B. E., Huang, H., McGarvey, P., Mazumder, R., & Wu, C. H. (2007). UNIREF: Comprehensive and non-redundant Uniprot Reference Clusters. Bioinformatics, 23(10), 1282– 1288. doi10.1093/bioinformatics/btm098

48. Richardson, L., Allen, B., Baldi, G., Beracochea, M., Bileschi, M. L., Burdett, T., Burgin, J., Caballero-Pe rez, J., Cochrane, G., Colwell, L. J., Curtis, T., Escobar-Zepeda, A., Gurbich, T. A., Kale, V., Korobeynikov, A., Raj, S., Rogers, A. B., Sakharova, E., Sanchez, S., … Finn, R. D. (2022). MGnify: The microbiome sequence data analysis resource in 2023. Nucleic Acids Research, 51(D1). doi10.1093/nar/gkac1080

49. Steinegger, M., Mirdita, M., & So ding, J. (2019). Protein-level assembly increases protein sequence recovery from metagenomic samples manyfold. Nature Methods, 16(7), 603–606. doi10.1038/s41592-019-0437-4

50. Steinegger, M., & So ding, J. (2018). Clustering huge protein sequence sets in linear time. Nature Communications, 9(1). doi10.1038/s41467-018-04964-5

51. Remmert, M., Biegert, A., Hauser, A., & So ding, J. (2011). HHblits: Lightning-fast iterative protein sequence searching by HMM-HMM alignment. Nature Methods, 9(2), 173–175. doi10.1038/nmeth.1818

52. *Introduction to Protein Data Bank Format*. Introduction to protein data bank format. www.cgl.ucsf.edu/chimera/docs/UsersGuide/tutorials/pdbintro.html

53. Jha, K., & Saha, S. (2020). Amalgamation of 3D structure and sequence information for protein–protein interaction prediction. Scientific Reports, 10(1). doi10.1038/s41598-020-75467-x

54. Rodrigues, C. H., Pires, D. E., Blundell, T. L., & Ascher, D. B. (2022). Structural landscapes of PPI interfaces. Briefings in Bioinformatics, 23(4). doi10.1093/bib/bbac165

55. Varadi, M., Anyango, S., Deshpande, M., Nair, S., Natassia, C., Yordanova, G., Yuan, D., Stroe, O., Wood, G., Laydon, A., Žídek, A., Green, T., Tunyasuvunakool, K., Petersen, S., Jumper, J., Clancy, E., Green, R., Vora, A., Lutfi, M., … Velankar, S. (2021). Alphafold protein structure database: Massively expanding the structural coverage of protein-sequence space with high-accuracy models. Nucleic Acids Research, 50(D1). doi10.1093/nar/gkab1061

56. Oates, M. E., Romero, P., Ishida, T., Ghalwash, M., Mizianty, M. J., Xue, B., Doszta nyi, Z., Uversky, V. N., Obradovic, Z., Kurgan, L., Dunker, A. K., & Gough, J. (2012). D2P2: Database of disordered protein predictions. Nucleic Acids Research, 41(D1). doi10.1093/nar/gks1226

57. Tunyasuvunakool, K., Adler, J., Wu, Z., Green, T., Zielinski, M., Žídek, A., Bridgland, A., Cowie, A., Meyer, C., Laydon, A., Velankar, S., Kleywegt, G. J., Bateman, A., Evans, R., Pritzel, A., Figurnov, M., Ronneberger, O., Bates, R., Kohl, S. A., … Hassabis, D. (2021). Highly accurate protein structure prediction for the human Proteome. Nature, 596(7873), 590–596. doi10.1038/s41586-021-03828-1

58. Adhikari, B., & Cheng, J. (2016). Protein residue contacts and prediction methods. Methods in Molecular Biology, 463–476. 10.1007/978-1-4939-3572-7_24

59. Slater, J. C. (1964). Atomic radii in Crystals. The Journal of Chemical Physics, 41(10), 3199– 3204. doi10.1063/1.1725697

60. Qing, R., Hao, S., Smorodina, E., Jin, D., Zalevsky, A., & Zhang, S. (2022). Protein design: From the aspect of water solubility and stability. Chemical Reviews, 122(18), 14085–14179. doi10.1021/acs.chemrev.1c00757

61. Graham, B., & van der Maaten, L. (2017, June 5). Submanifold sparse convolutional networks. CVPR*2018*

62. He, T., Zhang, Z., Zhang, H., Zhang, Z., Xie, J., & Li, M. (2018, December 5). Bag of tricks for image classification with Convolutional Neural Networks. CVPR2019

63. Huang, G., Liu, S., van der Maaten, L., & Weinberger, K. Q. (2018, June 7). CondenseNet: An efficient DenseNet using learned group convolutions. CVPR*2018*

64. Li, X., Han, P., Chen, W., Gao, C., Wang, S., Song, T., Niu, M., & Rodriguez-Pato n, A. (2022). MARPPI: Boosting prediction of protein–protein interactions with multi-scale architecture residual network. Briefings in Bioinformatics, 24(1). doi10.1093/bib/bbac524

65. Jing, X., Zeng, H., Wang, S., & Xu, J. (2019). A web-based protocol for Interprotein contact prediction by Deep Learning. Methods in Molecular Biology, 67–80. doi10.1007/978-1-4939-9873-9_6

66. Salam, A. (1991). The role of chirality in the origin of life. Journal of Molecular Evolution, 33(2), 105–113. doi10.1007/bf02193624

67. Jihong, J. Keras-resnet3d: Implementations of ResNets for volumetric data, including a vanilla resnet in 3D. GitHub. github.com/JihongJu/keras-resnet3d

68. Dudovitch, G. DENSENETFCN-3D: A 3D implementation of DenseNet & DenseNetFCN. GitHub. github.com/GalDude33/DenseNetFCN-3D

